# Impact of angiotensin-receptor blockers on intrarenal renin-angiotensin system activity in hypertension: A PK/PD modelling study

**DOI:** 10.1101/2023.07.13.548848

**Authors:** Delaney Smith, Anita Layton

**Affiliations:** Department of Applied Mathematics, University of Waterloo, 200 University Ave, Waterloo, N2L 3G1, ON, Canada; Cheriton School of Computer Science, University of Waterloo, 200 University Ave, Waterloo, N2L 3G1, ON, Canada; Department of Biology, University of Waterloo, 200 University Ave, Waterloo, N2L 3G1, ON, Canada; School of Pharmacy, University of Waterloo, 200 University Ave, Waterloo, N2L 3G1, ON, Canada

**Keywords:** renin-angiotensin system, angiotensin receptor blocker, losartan, hypertension, kidney, angiotensin II infusion, blood pressure regulation, PK/PD

## Abstract

The renin-angiotensin system (RAS) is a primary regulator of volume homeostasis and blood pressure, whose over-activation is commonly associated with hypertension. Indeed, medications that target the RAS are generally effective in reducing blood pressure. However, more can be learned about how these medications influence the intrarenal RAS. Angiotensin-receptor blockers (ARBs) in particular have been shown to exert different effects on the intrarenal and systemic RASs in various experimental models of hypertension. In rats chronically infused with angiotensin II (Ang II), ARBs consistently prevent intrarenal, but not systemic Ang II levels from rising. The former effect is sufficient in preventing the development of hypertension. The regulation of intrarenal RAS, independently of the systemic RAS, by ARBs has been hypothesized to be mediated by the inhibition of all positive feedback loops inherent to the intrarenal RAS, also known as the “key point breakdown effect.” To investigate the validity of this hypothesis, we developed a PK/PD model of the ARB Losartan that considers the kidney, and applied the model to study how this class of medication influences intrarenal RAS activity and consequently blood pressure regulation in male rats. Simulations indicate that ARBs more effectively inhibit the activation of the intrarenal RAS because, unlike in the plasma, this process relies on the accumulation of cell-associated Ang II. We hypothesize that it is by blocking this intracellular uptake pathway, and restricting Ang II to extracellular regions of the kidney where the peptide cannot initiate downstream signalling, that Losartan normalizes blood pressure. While the key point break down effect assists in this response, it alone is not sufficient. Our results highlight the intrarenal RAS as the key pharmacological target of ARB treatment and emphasize the importance of this local tissue RAS in the development of hypertension.

## 1 Introduction

Hypertension is the leading risk factor for cardiovascular mortality globally (Virani et al, 2020). Although the underlying causes of most cases of hypertension are unknown and likely multi-factorial, anti-hypertensive therapies that target the renin-angiotensin system (RAS) are common and effectual (Ibrahim, 2006), because of the long-established role of the RAS in blood pressure regulation (Fyhrquist et al, 1995; Sparks et al, 2014). Upon binding to angiotensin type 1 receptors (AT1Rs), angiotensin II (Ang II), the primary bio-active product of the RAS, increases blood pressure by inducing vasoconstriction, increasing renal sympathetic nervous activity, and stimulating sodium reabsorption (Fyhrquist et al, 1995; Sparks et al, 2014). Given that many of these actions take place within the kidney, an organ which has been found to not only express, but independently regulate all components of the RAS (Prieto-Carrasquero et al, 2004; Navar et al, 2011; Gonzalez and Prieto, 2015; Gonzalez et al, 2011; Navar et al, 2003; Kobori et al, 2001; Gonzalez-Villalobos et al, 2008; Zhuo et al, 2002, 1999; Cheng et al, 1995; Nishiyama and Kobori, 2018), the significance of the local intrarenal RAS in the pathogenesis and progression of hypertension has recently come into focus.

The systemic and intrarenal RAS are connected and typically vary in tandem. Indeed, in normal physiological states, the two systems are balanced by the filtration of peptides into and the flow of reabsorbed peptides out of the kidney. However, a decoupling of the two RAS has been observed in various experimental models of hypertension, including: Ang II-induced hypertensive rats (Zou et al, 1996), two-kidney, one-clip Goldblatt hypertensive rats (Cervenka et al, 1999), salt-sensitive rats (Wu et al, 2014), and spontaneously hypertensive rats (Takenaka et al, 1990). In Ang II-induced hypertension in particular, there is a progressive rise in intrarenal Ang II that cannot be explained on the basis of equilibration with plasma Ang II concentrations ([Ang II]) (Zou et al, 1996; Ellis et al, 2012; Gonzalez-Villalobos et al, 2008; Li et al, 2019). This response is prevented by treatment with angiotensin-receptor blockers (ARBs), a common anti-hypertensive medication that targets the RAS by binding to and blocking AT1Rs. *How does this class of medication influence the activity of the intrarenal RAS?* Addressing this question would significantly improve our understanding of clinical hypertension, and that can have wide benefits, given the commonality of intrarenal RAS over-activation across pre-clinical hypertensive models and the effectiveness of ARBs as an anti-hypertensive treatment strategy.

To explain how ARBs regulate intrarerenal RAS, it has been hypothesized that ARBs inhibit all positive feedback loops inherent to the intrarenal RAS, including the up-regulation of proximal tubule angiotensinogen (AGT), collecting duct renin, and tubular epithelial cell AT1R expression. This is also known as the “key point break down effect” (Xu et al, 2009). An objective of this study is to investigate the validity of this hypothesis. This is accomplished by coupling a model of the rat intrarenal RAS that incorporates the aforementioned feedback loops Smith and Layton (2023) to a pharmacokinetic (PK) model of an ARB to create a comprehensive pharmacokinetic/pharmacodynamic (PK/PD) model. In particular, we simulate and compare Ang II infusion experiments with and without ARB treatment to investigate two clinically relevant questions: (i) *Which Ang II accumulation mechanism, enhanced AT1R-mediated uptake (UPTK) or increased endogenous production (PROD), is the primary target of ARB treatment?* (ii) *How does this influence the general activity of the intrarenal RAS, and therefore blood pressure regulation?*

To accomplish these goals, we simulate the Ang II infusion experiments of Zou et al (1996), which were performed separately with and without continuous treatment with Losartan. Losartan was the first ARB prescribed to treat clinical hypertension, and remains an important drug in basic and clinical research today (Xu et al, 2009). Following its oral administration, Losartan is reabsorbed from gastrointestinal tract into the systemic circulation where, after entering the liver, it becomes metabolized into EXP3174 by cytochrome p450 enzymes (Tamaki et al, 1997). Both Losartan and its metabolite EXP3174 are competitive antagonists that selectively bind to AT1Rs with a high affinity (Miura et al, 2011; Xu et al, 2009).

In published computational models, the effect of Losartan on the RAS is often represented implicitly via a parameter that reduces Ang II-AT1R binding by an arbitrary target amount (Leete et al, 2018). However, a similar approach is not appropriate when we want the *predict* the effect of a specific ARB, based on the amount that has been administered. This is because the relationship between drug dosage and Ang II-AT1R binding is rarely, if ever, explicitly experimentally quantified. Instead, a PK model describing Losartan and its metabolite EXP3174 is desirable, as it can be coupled to our previous RAS model (Smith and Layton, 2023) via the drug’s known AT1R binding properties. Such a model has previously been developed to study gastric emptying in humans (Karatza and Karalis, 2020). However, that model simulates a single oral dose of the drug (not continuous treatment with Losartan), and does not consider the kidney as a separate compartment (thus not appropriate for studying intrarenal RAS).

In this work, we develop a PK/PD model of the ARB Losartan that considers the kidney and simulate the Ang II and ARB administration experiments performed by Zou et al (1996). Simulation results elucidate the mechanisms by which ARBs effectively target the intrarenal RAS to prevent its dis-regulation and consequently, the development of hypertension following Ang II infusion.

## 2 Methods

### 2.1 RAS model

The RAS model referenced in this work is taken directly from Smith and Layton (2023). It considers both the systemic and the intrarenal RAS via a systemic compartment and four intrarenal tissue compartments: the glomerular compartment, the peritubular compartment, the tubular compartment, and the vasculature compartment. The renal vasculature compartment comprises of both the lymphatic and the post-glomerular blood vasculature. The preglomerular blood vasculature is considered part of the systemic compartment. The other renal compartments are sub-divided into intracellular and extracellular regions, connected to one another via megalin- and AT1R-dependent Ang II internalization. For a complete description of the RAS model and how it may be used to simulate Ang II infusion experiments, see Smith and Layton (2023).

### 2.2 Losartan-EXP3174 pharmacokinetic model

The PK model developed in this work considers Losartan and EXP3174 dynamics in the rat across four compartments: the gastrointestinal (GI) tissue compartment, the systemic vasculature compartment, the peripheral tissue compartment, and the renal tissue compartment. Orally-administered Losartan first enters the GI compartment, before being absorbed into the systemic blood plasma (the systemic compartment) where a portion is converted into its bio-active metabolite EXP3174 by cytochrome p450 enzyme activity (Tamaki et al, 1997). Both Losartan and its metabolite are then re-distributed into the poorly perfused organs (the peripheral tissue compartment) and the kidney (the renal compartment). The renal compartment is divided into the same tissue sub-compartments as the intrarenal RAS model outlined in our previous work (Smith and Layton, 2023) and summarized in section 2.1. The PK model is coupled to that of the intrarenal RAS through Losartan and EXP3174-AT1R-binding in the systemic and intrarenal compartments (see section 2.3). The equations used to model the concentrations of Losartan and EXP3174 in each compartment are described in detail below, with a schematic diagram provided in Figure I.

**Fig. I.**
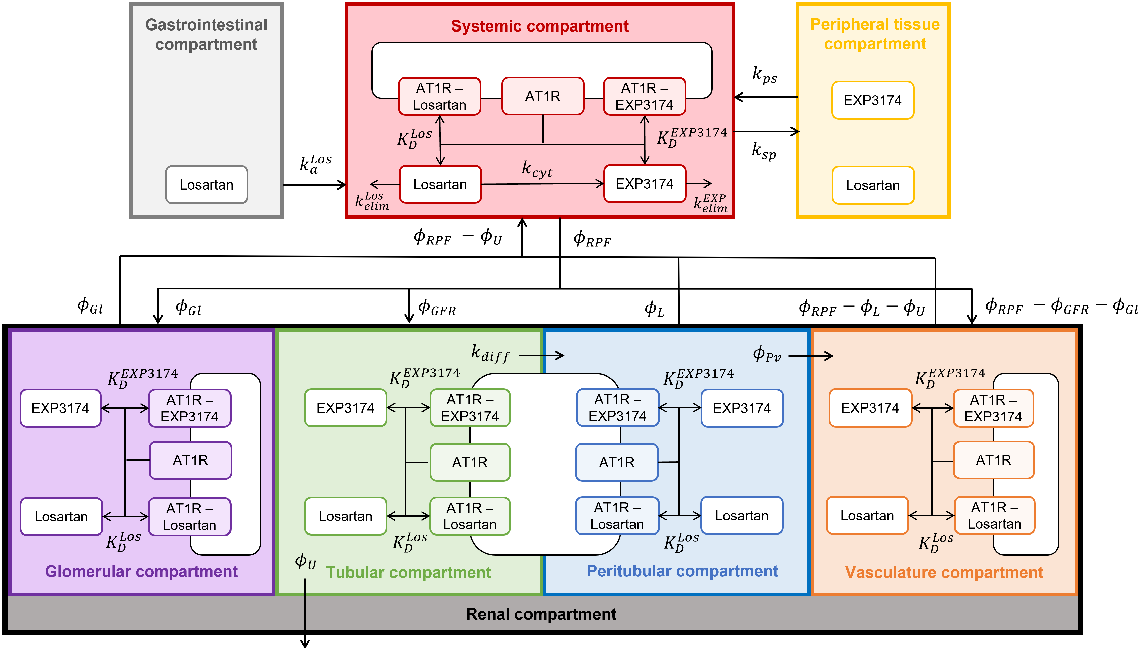
Schematic representation of the Losartan-EXP3174 PK model. The definition of all variables can be found in the text. Orally-administered Losartan enters the circulation (systemic compartment; red) via absorption from the gastrointestinal compartment (light grey) where a portion is converted into its bio-active metabolite EXP3174 by cytochrome p450 enzymes. These molecules are then re-distributed into the poorly perfused organs (the peripheral tissue compartment; yellow) and the kidney (the renal compartment; black). In the kidney, the molecules may either bind to regional AT1Rs, be excreted in urine (at a rate of *ϕ*_*U*_), or get reabsorbed back into the systemic compartment.

#### 2.2.1 Gastrointestinal compartment

The rate of change of the amount of Losartan in the GI compartment, *Los*_*GI*_, is determined by the balance between the drug’s administration rate *K*_*Los*_ and its absorption into the systemic compartment with rate constant 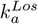:

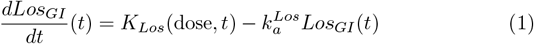

We simulate two different modes of drug delivery, via a single oral dose and in drinking water. The respective simulations differ in their definition of the source term *K*_*Los*_(dose, *t*) and the initial amount *Los*_*GI*_ (0). To simulate a single oral dose *D* (fmol) of Losartan, we set

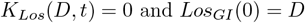

Thus, the given dose of Losartan is absorbed into the systemic compartment upon being administered at *t* = 0. To simulate a dose *D* (fmol) of Losartan administered in drinking water each day, we set

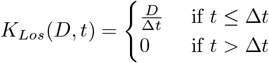

where ∆*t* = 12 hours and *Los*_*GI*_ (0) = 0. Indeed, the rat is assumed to be awake, constantly drinking water during the first 12 hours of the day and asleep for the remaining 12 hours. In this way, they consume their entire daily dose of Losartan in their awake period. To simulate a drug administered in drinking water over multiple days, we run the model iteratively, using the last time point of the previous iteration as the initial condition for the next iteration.

#### 2.2.2. Systemic vasculature compartment

Losartan begins accumulating in the systemic plasma upon being reabsorbed from the GI compartment with rate 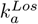. In plasma, Losartan can be converted to EXP3174 by cytochrome p450 enzymes (Tamaki et al, 1997) with rate *k*_*cyt*_, degraded naturally with rate 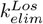, redistributed to the peripheral compartment with rate *k*_*sp*_, or filtered into the kidney (the renal compartment) at a rate proportional to renal plasma flow *ϕ*_*RPF*_ (scaled by the ratio of kidney weight *W*_*K*_ to circulating plasma volume *V*_*circ*_). Losartan subsequently returns to the systemic circulation from the peripheral compartment at rate *k*_*ps*_, the peritubular interstitial space at a rate proportional to lymphatic flow *ϕ*_*L*_, and the renal (post-glomerular) blood vasculature at a rate proportional to *ϕ*_*RPF*_ – *ϕ*_*L*_ – *ϕ*_*U*_. Finally, plasma Losartan may to bind to and unbind from systemic AT1Rs (Eq. 7) with rates 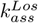 and 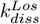, respectively.

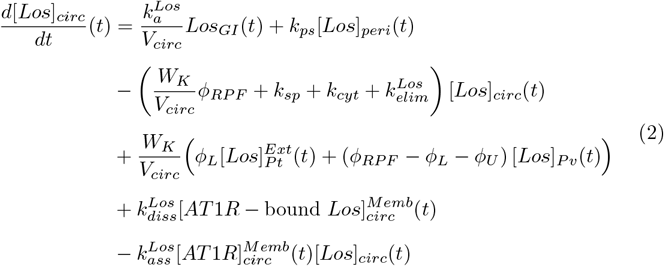

In the systemic plasma, EXP3174 is produced from Losartan via cytochrome p450 enzyme activity with rate *k*_*cyt*_. Otherwise EXP3174 behaves identically to Losartan in this compartment, differing only in the rate of elimination 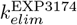, AT1R binding 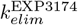, and AT1R unbinding 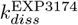.

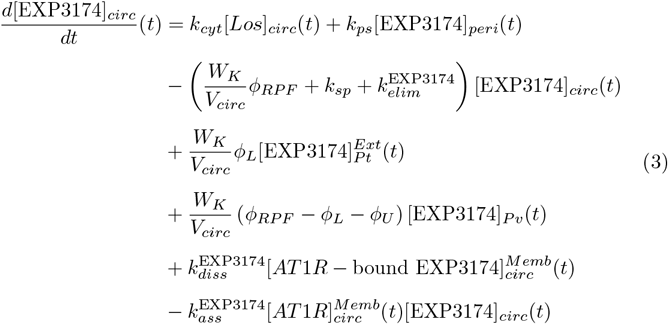

#### 2.2.3 Peripheral tissue compartment

Both Losartan and EXP3174 are assumed to enter and exit the peripheral compartment at the same rates of *k*_*sp*_ and *k*_*ps*_, respectively. Consequently the rate of change of the peripheral concentration of metabolite *X* (*X* = *Los* or *X* = EXP3174) is given by:

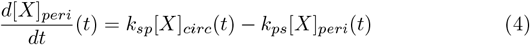

#### 2.2.4 Renal tissue compartment

In the kidney, the concentration of Losartan and EXP3174 is described by their redistribution among the extracellular regions of the intrarenal subcompartments and their association/dissociation with local membrane-bound AT1Rs. The structure of the equations describing the dynamics of each metabolite is similar to that of Ang I and II in the intrarenal RAS model and as such, is similar across all intrarenal sub-compartments. Therefore, as done in Smith and Layton (2023), we introduce a general equation describing the rate of change of extracellular Losartan (*X* = *Los*) and EXP3174 (*X* = *EXP* 3174) in compartment *C* below, where *C* = *Gl, Tb, Pt*, or *Pv*.

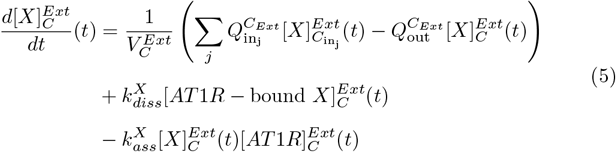

The first line represents the balance between the *j* fluxes into subcompartment *C*_*Ext*_ from sub-compartment 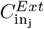 with rate 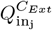 and the flux out of sub-compartment *C*_*Ext*_ with rate 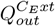. The number of incoming fluxes *j* differs for each compartment, as summarized in Table I. 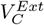 is a parameter describing the volume of each extracellular sub-compartment. All flux rates and volume parameters are consistent with those of the intrarenal RAS model (Smith and Layton (2023); Table AI). The second last and last lines of Eq. 5 represents the unbinding and binding of metabolite *X* to membrane-bound AT1Rs with rates 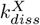 and 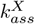, respectively.

**Table I.** Intrarenal sub-compartment-specific parameters corresponding to Eq. 5.

| **C** | $Q_{inj}^{C_{Ext}}$ | $[X]_{C_{inj}}^{Ext}(t)$ | $Q_{out}^{C_{Ext}}$ |
| --- | --- | --- | --- |
| **Gl** | $\phi_L$ | $[X]_{circ}(t)$ | $\phi_L$ |
| **Tb** | $\phi_{GFR}$<br>$\phi_{Gl}$ | $[X]_{circ}(t)$<br>$[X]_{Gl}^{Ext}(t)$ | $k_{diff} + \phi_U$ |
| **Pt** | $k_{diff}$ | $[X]_{Tb}^{Ext}(t)$ | $\phi_{Pv} + \phi_L$ |
| **Pv** | $\phi_{RPF} - \phi_{GFR} - \phi_{Gl}$<br>$\phi_{Pv}$ | $[X]_{circ}(t)$<br>$[X]_{Pt}^{Ext}(t)$ | $\phi_{RPF} - \phi_L - \phi_U$ |

For the complete set of Losartan-EXP3174 model equations in the renal compartment, see Appendix A. For a detailed schematic of the Losartan-EXP3174 model, see Figure I.

### 2.3 Coupling the RAS and Losartan-EXP3174 models

As aforementioned, the Losartan-EXP3174 model is coupled to that of the RAS through Losartan- and EXP3174-AT1R binding in the systemic and renal model compartments. We assume for simplicity that membrane-bound AT1R-Losartan and AT1R-EXP3174 complexes are not internalized. In this way, the dynamics of these complexes are described by substrate-receptor association and dissociation with rates 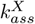 and 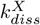, respectively, where *X* = *Los* or EXP3174:

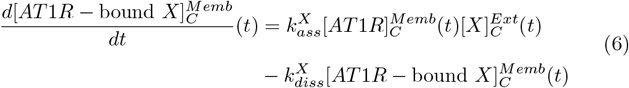

In this general equation, *C* = *circ, Gl, Pt, Tb*, or *Pv*. To account for AT1R binding to Losartan and EXP3174 in the RAS model, we update the previous equations describing membrane-bound free AT1R dynamics in all compartments as follows:

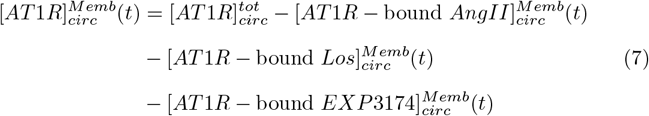

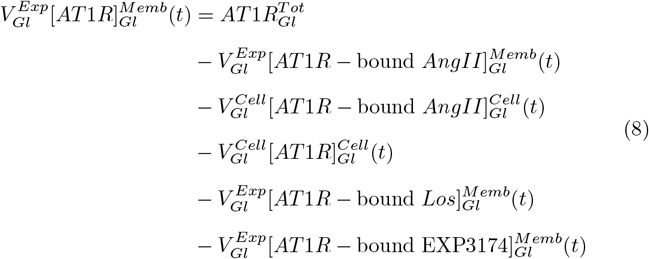

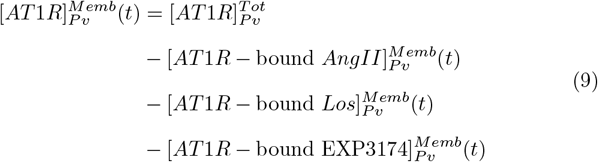

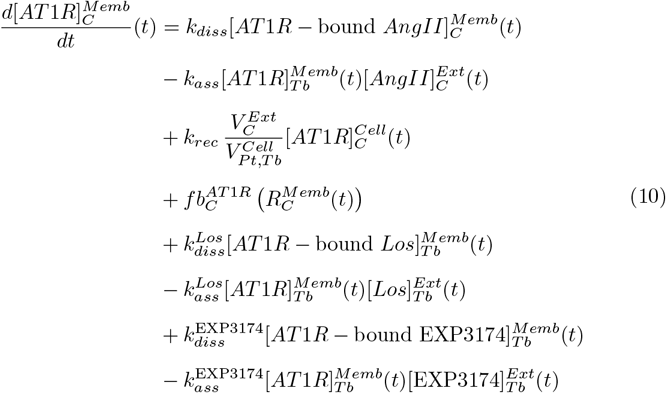

where *C* = *Tb* or *Pt*.

To facilitate running *in silico* Losartan administration experiments, we also modify the secretion of renin from the juxtamedullary apparatus such that the strength (*B*_*AT* 1*R*_; Eq. 13) of the AT1R- and Ang II-dependent feedback (*ν*_*AT* 1*R*_; Eq. 12) can respond in a situation where renin is meant to increase or decrease, i.e. when AT1R-bound Ang II decreases or increases, respectively. In the previous formulation (Smith and Layton, 2023), the fitting of *B*_*AT* 1*R*_ was restricted to a situation where renin secretion was meant to decrease, i.e. Ang II infusion experiments. In contrast, following Losartan administration renin secretion is expected to increase. Hence, we allow *B*_*AT* 1*R*_ to depend on the ratio of 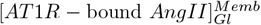 to control 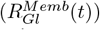. Indeed, plasma renin concentration is modelled via using the following expression:

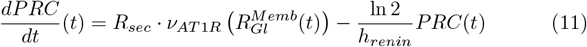

where

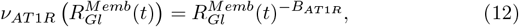

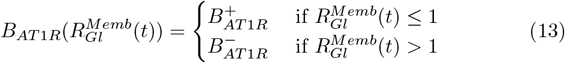

We take 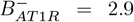, the value previously fit to Ang II infusion data by Smith and Layton (2023) and determine 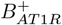 using Losartan administration data (section 2.4).

Finally, we maintain the assumption that renal renin activity 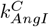 remains constant following Losartan administration. This has been observed experimentally in the case of Ang II infusion Shao et al (2009). However, to our knowledge, similar data is not available following Losartan treatment. We therefore base this assumption on the results presented in Smith and Layton (2023), that implicit positive feedback on intrarenal AGT and renin production counteracts the decrease in filtered renin from the systemic circulation. Given this, it seems plausible that any increase in filtered renin (caused by an increase in PRC following Losartan treatment) is also balanced by the inhibition of local intrarenal AGT and renin production.

### 2.4 Model parameter identification

The compartment volume and renal hemodynamic parameters present in the PK model were taken directly from the RAS model’s parameter set (Smith and Layton (2023); Table AI). It was assumed that the rate of association of AT1Rs with Losartan and EXP3174 was the same as Ang II, i.e. that 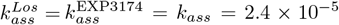 /fmol/mL per min (Schalekamp and Danser, 2006). The dissociation rate for each metabolite *X* (*X* = *Los*, EXP3174) was then computed using the expression 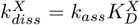, where 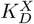 is the dissociation constant of metabolite *X* reported in the literature (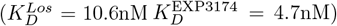) (Miura et al, 2011). The remaining model parameters (apart from 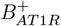; Eq. 13) were estimated by simulating a single oral dose (10 mg/kg) of Losartan and minimizing the sum of squared errors between the simulated and experimental plasma drug concentration time series from Li et al (2016). Lastly, 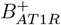 was identified by administering 30 mg/kg of Losartan per day in drinking water to the model and comparing the average fold-change in PRA on day 13 to the data from Zou et al (1996).

## 3 Results

### 3.1 Model parameter identification

**Table II.**
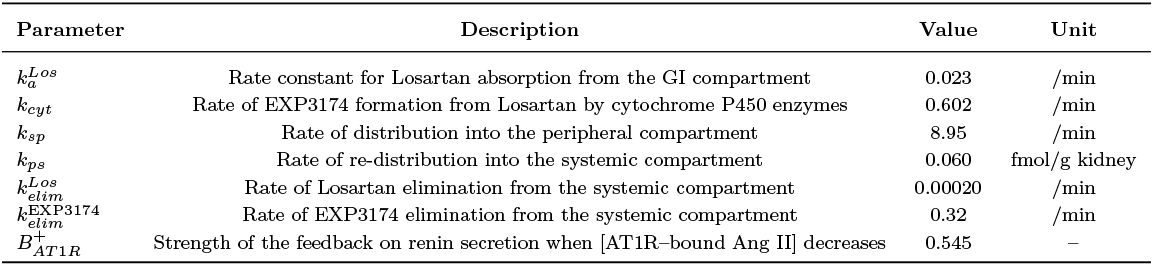
Losartan-EXP3174 model parameters.

As outlined in section 2.4, a single oral dose (10 mg/kg) of Losartan was administered to the model and the error between the true and simulated plasma [Losartan] and plasma [EXP3174] time series was minimized to identify the model parameters. The optimized parameter set is given in Table II, with the simulated (curves) and true time series (markers) shown in Figure II.

**Fig. II.**
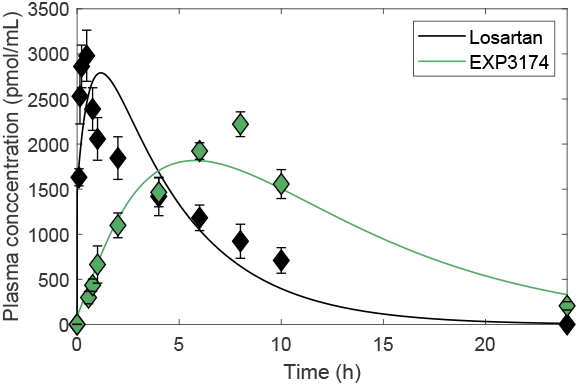
Model fit (curves) to plasma Losartan and EXP3174 concentration time series data (markers) following a single oral dose (10 mg/kg) of Losartan.

Below, fitting results specific to the strength of Ang II-AT1R feedback on renin secretion 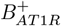 are detailed and the model is validated against the remaining Ang II infusion and Losartan administration experimental data from Zou et al (1996).

### 3.2 Model validation

In addition to the Ang II infusion experiment (experiment (i): *Ang II*) replicated in Smith and Layton (2023), Zou et al (1996) also performed two Losartan administration experiments, where 30 mg/kg of Losartan was administered per day in drinking water either alone (experiment (ii): *Losartan*) or in conjunction with 40 ng/min of Ang II (experiment (iii): *Ang II + Losartan*). Replicating these experiments *in silico* forms an important step towards the validation of our model.

Figure III compares the results of each simulated experiment to the data that was collected by Zou et al (1996) on day 13. In particular, the average (day 13) simulated (wide bars) and experimental (narrow bars) RAS peptide concentrations are compared to their control values (i.e. Smith and Layton (2023)’s model steady state; Table III). The results of experiment (i) have already been detailed previously in Smith and Layton (2023) and are therefore shown for comparison purposes. All data points relating to experiments (ii) and (iii), apart from the increase in PRA under Losartan administration alone (Figure III, starred bar) which was used to fit 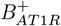 (see section 2.4), were kept for model validation. In general, the simulations show good agreement to data, capturing the experimental trends well. This not only speaks to the robustness of the PK/PD model, whose PK parameters were identified using data obtained via a different drug administration route (single oral dose vs. in drinking water) and over a different time scale (1 day vs. 2 weeks), but also to the adaptability of the original RAS model, whose parameters (apart from the addition of 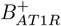 did not need to be modified to accurately predict the variations in systemic and intrarenal RAS peptides that are induced by Losartan.

**Fig. III.**
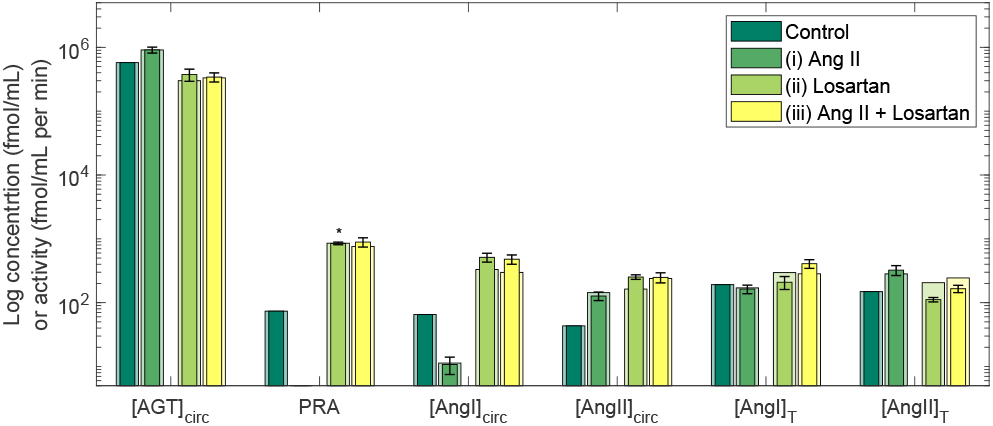
Model validation (wide bars) against RAS peptide data (experimental) on day 13 of experiment (i): 40 ng/min subcutaneous Ang II infusion; experiment (ii): 30 kg/mg/day Losartan administration in drinking water; and experiment (iii): 40 ng/min subcutaneous Ang II infusion alongside 30 kg/mg/day Losartan administration in drinking water of Zou et al (1996). Control (steady state) concentrations are also provided for comparison.

Model simulations indicate that once administered, Losartan and EXP3174 rapidly bind to systemic and intrarenal AT1Rs, significantly reducing the concentration of AT1R–bound Ang II in all compartments (see section 3.3). With 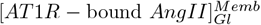 significantly reduced, the secretion of renin from the juxtamedullary apparatus of the kidney is increased (Eq. 12) causing PRA to rise significantly. As a result, plasma [AGT] decreases and plasma [Ang I] increases above control. Model simulations confirm that the observed decrease in plasma [AGT] is the consequence of both enhanced renin activity and a blockage of positive-feedback on hepatic AGT production, as was proposed by Ref. Zou et al (1996). The increase in plasma [Ang I] in conjunction with the unbinding of Ang II from systemic AT1Rs results in a similar fold-increase in plasma [Ang II]. As a result, more Ang I and Ang II get filtered into the kidney, causing the renal [Ang I] and [Ang II] to also increase slightly beyond control. All aforementioned changes in all RAS peptide levels following experiments (ii) and (iii) are summarized in Figure III.

In the next section, we examine the experimental results of Zou et al (1996) in more detail and use model simulations to offer further insight into the impacts of Losartan on both the systemic and intrarenal RAS and the inception of hypertension.

### 3.3 Model predictions

#### Losartan blocks the activation of all systemic and intrarenal positive feedback, resulting in similar regulation of the RAS in the absence and presence of Ang II infusion

In their study, Zou et al (1996) showed that each RAS peptide varies similarly regardless of whether Ang II is infused concurrently with Losartan or not. In other words, the peptide levels following experiment (ii) and experiment (iii) were similar (Figure III). Model simulations indicate that this is because Losartan inhibits all systemic and intrarenal positive feedback loops by blocking AT1R-Ang II binding: With AT1Rs blocked by Losartan, excess Ang II from the infusion cannot initiate any downstream signalling, including the activation of all AT1R-mediated positive feedback, and thus the intrinsic regulation of all endogenous RAS peptides remains approximately the same as if an infusion were not taking place. As a result, the only peptide whose concentration significantly differs following Ang II infusion under concurrent Losartan administration is Ang II itself. Indeed, by separating the endogenously produced Ang II from the exogenously infused Ang II (for details see Smith and Layton (2023), section 2.3.2), we show that exogenous Ang II is the primary contributor to the increased plasma [Ang II] (Figure IVa) and intrarenal [Ang II] (Figure IVb) observed in experiment (iii) relative to experiment (ii). In particular, the overall fold-increase in plasma [Ang II] is larger than the fold-increase in intrarenal [Ang II] because the kidney accumulates comparatively less exogenous Ang II than the systemic compartment. The mechanisms governing this disparity are outlined below.

**Fig. IV.**
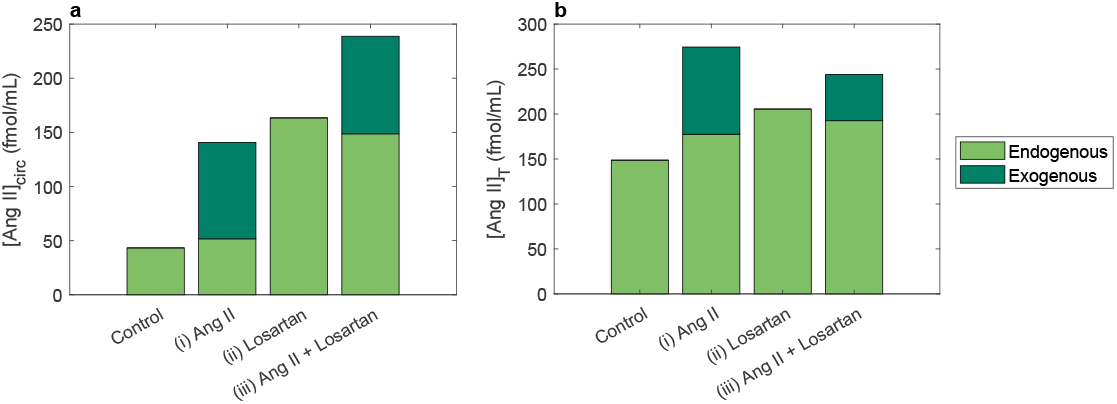
Average **a** plasma and **b** intrarenal exogenous (dark green) and endogenous (light green) [Ang II] on day 13 of experiment (i): 40 ng/min subcutaneous Ang II infusion; experiment (ii): 30 kg/mg/day Losartan administration in drinking water; and experiment (iii) 40 ng/min subcutaneous Ang II infusion alongside 30 kg/mg/day Losartan administration in drinking water of Zou et al (1996).

#### Losartan has differential effects on the accumulation of exogenous and endogenous Ang II in the plasma and the kidney, as a result of their independent regulation

As outlined in the previous Chapter, less exogenous Ang II accumulates in the kidney than in the plasma during a normal Ang II infusion (experiment (i)). However, this disparity is significantly enhanced by concurrent Losartan administration (experiment (iii)). Indeed, model simulations indicate that Losartan reduces the accumulation of exogenous Ang II in the kidney, but not the systemic circulation following Ang II infusion (Figure IV; bars (i) vs. (iii), dark green). In particular, its administration resulted in a 47% decrease in the simulated intrarenal exogenous [Ang II] with no change in the systemic exogenous [Ang II]. These results are qualitatively consistent with those of Zou et al (1998), who observed a marked decrease in the exogenous intrarenal, but not plasma, [Ang II] following concurrent Ang II infusion and Losartan administration. This also supports the hypothesis presented in Smith and Layton (2023), that intrarenal Ang II accumulation relies primarily on AT1R-mediated uptake of Ang II into cells during Ang II infusion. Because the accumulation of exogenous Ang II in the plasma does not rely on AT1Rs (occurs extracellularly), Losartan has little impact on this concentration. In contrast, the endogenous concentration of plasma Ang II, but not intrarenal Ang II, is greatly enhanced by Losartan following Ang II infusion (Figure IV; bars (i) vs. (iii), light green). This effect was also reported by Zou et al (1998), who observed little change in intrarenal endogenous [Ang II] alongside a significant increase in plasma endogenous [Ang II] when Losartan was administered concurrrently with Ang II. Model simulations indicate that the increase in endogenous plasma [Ang II] arises as a result of Losartan displacing all endogenous Ang II peptides that were initially bound to AT1Rs, and therefore not included in the free [Ang II]. Although this displacement also happens in the kidney, [*AngII*]_*T*_ depends on the concentration of AT1R-bound Ang II in all sub-compartments (Smith and Layton (2023); Eq. (13)), and therefore Losartan has a smaller impact on the endogenous intrarenal [Ang II].

#### Losartan likely prevents hypertension induced by Ang II infusion by restricting Ang II to regions where it cannot initiate downstream signalling

By examining the effects of Losartan on the intrarenal distribution of Ang II, we can gain insight into the mechanisms by which this anti-hypertensive therapy mitigates the consequences of sustained Ang II infusion. In particular, Figure V compares the temporal change in the intrarenal distribution of [Ang II] following Ang II infusion alone (panels x.i.) and in conjunction with Losartan administration (panels x.iii.). As outlined above, Losartan administration significantly reduces the amount of exogenous Ang II (dark green) that accumulates in the kidney during sustained Ang II infusion (Figure Va). It does this by rapidly binding to all AT1Rs within the kidney, displacing the previously AT1R-bound endogenous Ang II (light green) and preventing any subsequent AT1R-Ang II binding because of it’s (and EXP3174’s) high receptor affinity (Miura et al, 2011). Indeed, the membrane-bound fraction of intrarenal Ang II rapidly decreases to 0 within the first 5 hours of Losartan administration (Figure Vb). This blockage occurs before any exogenously infused Ang II even has a chance to bind to these receptors. As a result, Ang II no longer accumulates within the tubular epithelium (Figure Vc). In fact, the intracellular fraction of intrarenal Ang II decreases by approximately 69% following Losartan administration, from 0.35 to 0.11. This small intracellular fraction is entirely sustained by megalin-dependent uptake, which is unaffected by Losartan administration. With intracellular uptake severely limited, Ang II is forced to accumulate in the extracellular regions of the kidney (Figure Vd). The forced shift from intracellular to extracellular Ang II accumulation explains Losartan’s effectiveness as an anti-hypertensive therapy: Although the total intrarenal [Ang II] still increases slightly beyond control, Losartan importantly restricts Ang II to compartments where the peptide cannot activate any downstream signalling cascades, such as the stimulation of sodium reabsorption. As a result, blood pressure does not increase (Zou et al, 1996).

**Fig. V.**
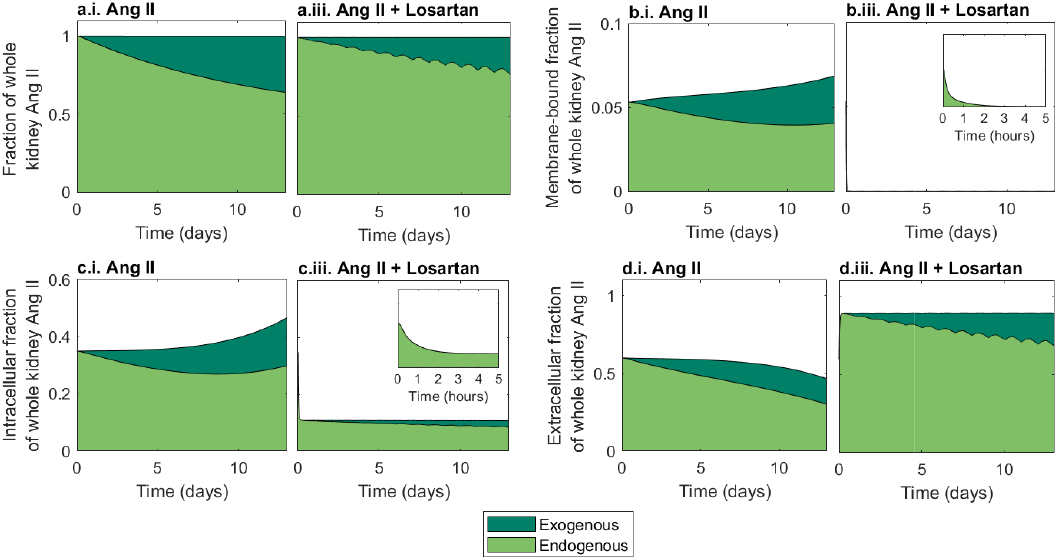
Temporal change in the intrarenal distribution of Ang II throughout 13 days of SC Ang II infusion (40 ng/min) alone (panels **x.i**) or in conjunction with 30mg/kg/day Losartan administration in drinking water (panels **x.iii**). Panels depict the relative contribution of endogenous (light green) and exogenous (dark green) Ang II to the: (**a**) total renal [Ang II], (**b**) membrane-bound fraction of renal [Ang II], (**c**) intracellular fraction of renal [Ang II], and (**d**) extracellular fraction of renal [Ang II].

In a real-world scenario, Losartan is prescribed when a patient has already developed hypertension, and not as a preventative measure. Therefore we use the model to simulate the effect of Losartan on a hypertensive rat, whose high blood pressure has been induced by a 40 ng/min 13-day SC Ang II infusion (experiment (i) from Zou et al (1996)). Figure VI illustrates the effect of Losartan treatment on the intrarenal concentration (panel a) and distribution (panels b-f) of Ang II. All observations are consistent with those of experiment (iii): Losartan inhibits the uptake of circulating (exogenous) Ang II within the kidney, but does not fully return intrarenal Ang II levels to control (Figure VIa, b). Nevertheless, Losartan drastically alters the intrarenal distribution of Ang II such that the vast majority becomes restricted to extracellular compartments (Figure VIc, d, e). Since the accumulated Ang II can no longer act as a signalling molecule in these regions, blood pressure is likely to decrease as a result of the Losartan administration.

**Fig. VI.**
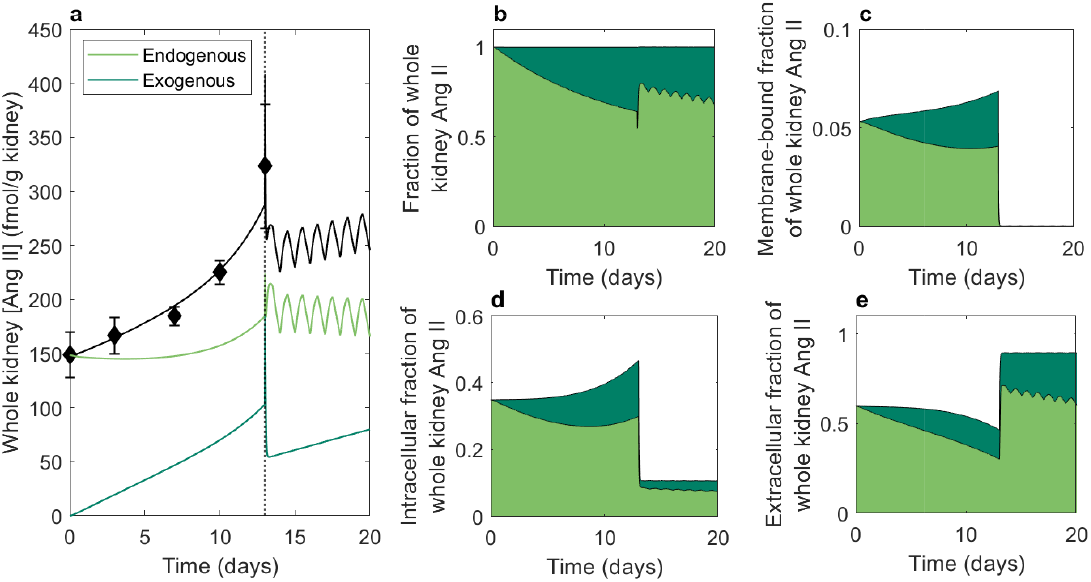
The effect of Losartan on the intrarenal (**a**) concentration and (**b**–**e**) distribution of Ang II in a hypertensive rat. Hypertension was induced via 13 days of subcutaneous Ang II infusion at 40 ng/min (experiment (i) from Zou et al (1996); diamonds) and subsequently treated with Losartan (30 mg/kg/day) for 7 days. Panels **b**–**e** depict the relative contribution of endogenous (light green) and exogenous (dark green) Ang II to the: (**b**) total renal [Ang II], (**c**) membrane-bound fraction of renal [Ang II], (**d**) intracellular fraction of renal [Ang II], and (**e**) extracellular fraction of renal [Ang II].

### 3.4 Local sensitivity analysis

To determine the robustness of the aforementioned results, a local parametric sensitivity analysis was performed after simulating experiment (iii) from Zou et al (1996): 40 ng/min subcutaneous Ang II infusion plus 30 mg/kg/day Losartan administration in drinking water. Indeed, the percent change in the average drug (Figure VII) and RAS peptide (Figure VIII) concentrations on the final day of the experiment (day 13) following a 10% increase in each model parameter was computed in each model compartment.

**Fig. VII.**
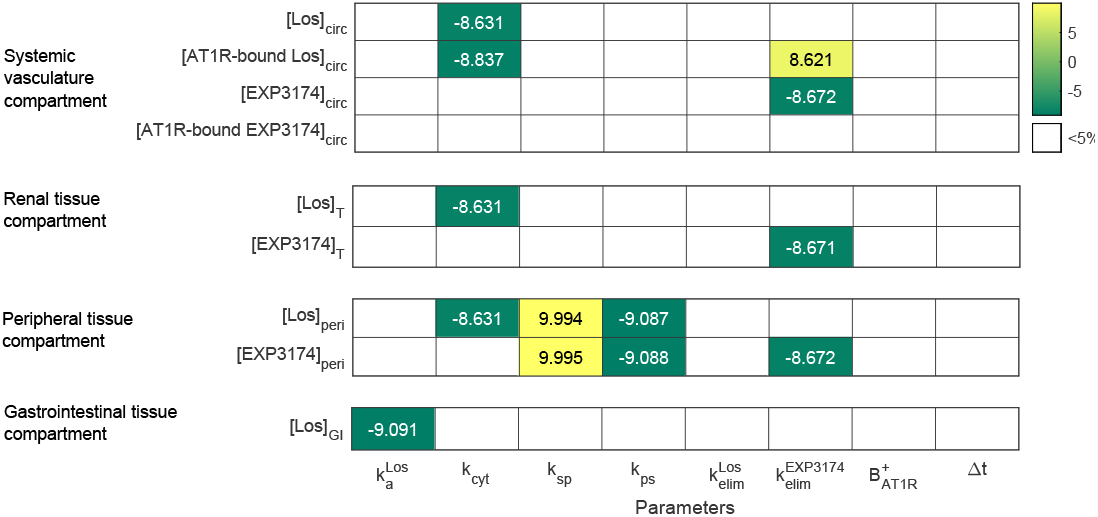
Percent change in the predicted average compartmental Losartan and EXP3174 concentrations following 13 days of 40 ng/min SC Ang II infusion alongside 30 mg/kg/day Losartan administration in drinking water when each parameter value is increased by 10%.

As shown in Figure VII, the effects of the parameteric perturbations on the concentrations of Losartan and EXP3174 in all compartments were as expected. When the Losartan dose is reabsorbed more rapidly from the gastrointestinal tract (i.e. 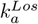 is increased), the drug’s average concentration in the GI tissue compartment on day 13 of treatment is lower than control. When the activity of systemic cytochrome p450 enzymes *k*_*cyt*_ is increased, and therefore the conversion of Losartan to EXP3174 is increased, the Losartan concentration in all downstream compartments (systemic, renal, and peripheral) decreases. An increased rate of uptake *k*_*sp*_ or release *k*_*ps*_ of Losartan from the peripheral tissue compartment results in a higher or lower Losartan concentration in these tissues, respectively. When EXP3174 is systemically degraded at a more rapid rate (i.e. 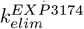 increases), it’s concentration in all downstream compartments decreases. Systemically, this frees up more AT1Rs for Losartan to bind to, thereby increasing the concentration of AT1R-bound Losartan in this vasculature compartment. Unlike EXP3174 however, increasing the rate of Losartan degradation 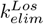 does not affect significantly affect the drugs concentration. Given this, and the fact that it’s optimized value is very close to 0, we can stipulate that Losartan degradation is primarily mediated by its conversion to EXP3174. Finally, increasing the drinking water duration ∆*t*, and therefore the time over which the rat consumes the drug dose does not impact it’s average concentration in any compartment as the amount of drug that is consumed overall remains unchanged. Moreover, the strength of the feedback on renin secretion 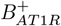 does not impact the PK model itself, only the intrarenal RAS model that it is coupled to.

In fact, as shown in Figure VIII, the intrarenal RAS model predictions are most sensitive to increases in 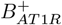. With the feedback loop’s strength increased, the same fold-decrease in AT1R-bound Ang II elicits a larger secretion of renin from the kidney and therefore a larger production of all endogenous peptides downstream in the cascade (Ang I and Ang II). As a result, more endogenous Ang II also gets filtered into the kidney which raises the total intrarenal [Ang II] in all regions. As expected, changes to 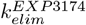 do not affect exogenous Ang II levels. The only other parameter that impacts intrarenal RAS model predictions is the rate of EXP3174 degradation in the systemic compartment 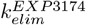. Since EXP3174 is much more potent than Losartan, it mediates the majority of its AT1R-blocking effects (Rossi, 2009). Therefore, when its degradation rate is increased (and therefore, its concentration is decreased), less AT1Rs are blocked by the metabolite, allowing for more AT1R-Ang II binding. However, the resulting increase in both systemic and intrarenal AT1R-bound Ang II, though proportionally large, are minute in absolute value (on the order of 10^*−*^3). Therefore, these changes do not influence the rest of the system significantly.

**Fig. VIII.**
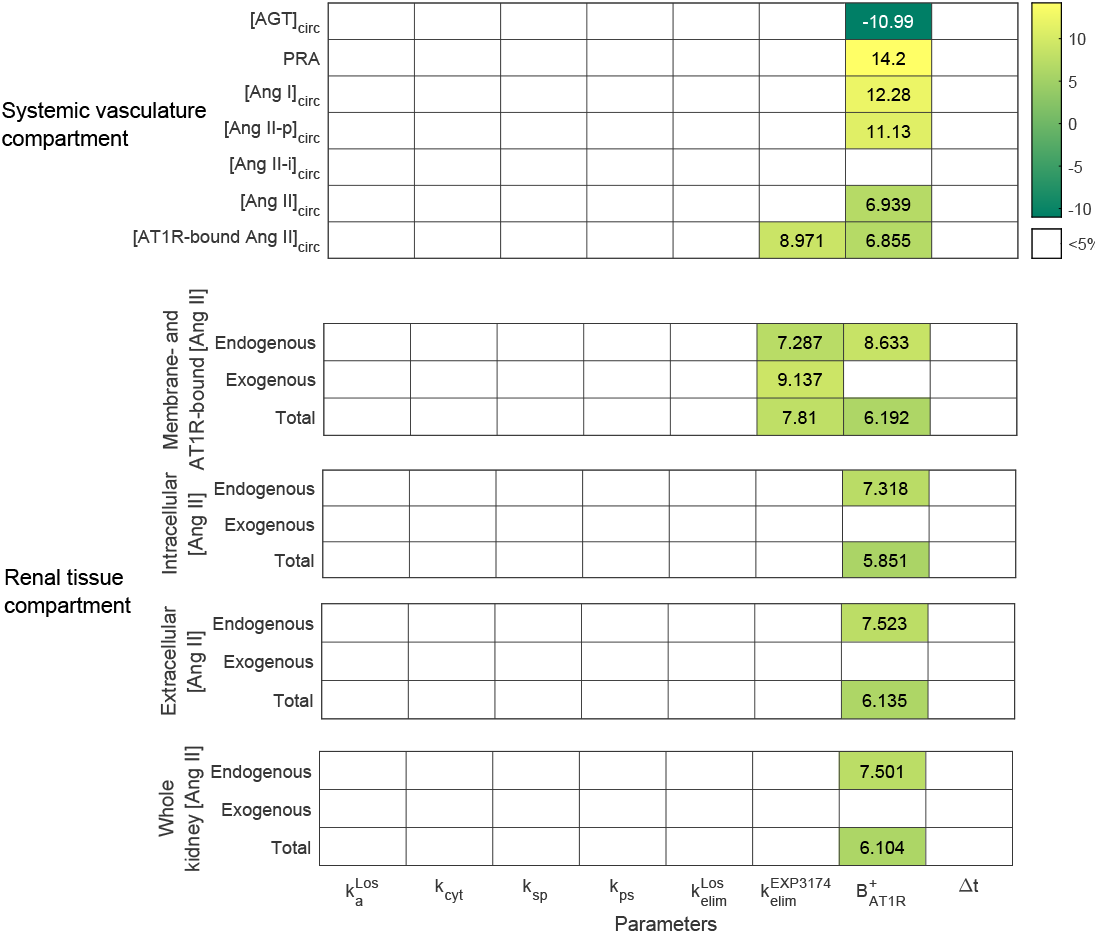
Percent change in the predicted average systemic and intrarenal RAS peptide concentrations following 13 days of 40 ng/min SC Ang II infusion alongside 30 mg/kg/day Losartan administration in drinking water when each parameter value is increased by 10%.

## 4 Discussion

In this work, we aimed to study the effects of antihypertensive therapy Losartan on the activity of the intrarenal RAS in both control and hypertensive conditions. To do so, a robust PK model of Losartan and its bioactive metabolite EXP3174 was developed and coupled to that of the intrarenal and systemic RAS presented in Smith and Layton (2023). The coupled model was then used to replicate the various Losartan administration experiments from Zou et al (1996). In particular, Losartan (30 mg/kg/day) was administered in drinking water either alone or in conjunction with sustained low dose (40 ng/min) Ang II infusion and the results were examined in detail.

Studies have shown that Losartan treatment helps to prevent intrarenal Ang II accumulation during Ang II infusion experiments and other experimental models of hypertension Zou et al (1996, 1998); Xu et al (2009). Model simulations indicate that this is mainly mediated by a reduced uptake of circulating (exogenous) Ang II into tubular epithelial cells. Endogenous peptide levels in the kidney remain relatively unchanged despite significantly increased plasma concentrations. We propose that this is due to Losartan’s inhibition of all positive feedback on endogenous Ang II production in the kidney (which is implicit in the assumption of a constant renal renin activity, see section 2.2; renal tissue compartment). These observations are qualitatively consistent with the results of Zou et al (1998), who reported decreased exogenous, but not endogenous intrarenal Ang II levels following sustained Losartan treatment. Indeed, it is likely that enhanced AT1R-mediated uptake, facilitated by increased AT1R expression, is the primary mechanism by which Ang II accumulates in the kidney during Ang II infusion. Positive feedback on local AGT and renin production acts secondarily to conserve basal Ang II production rates despite significantly reduced plasma, and therefore filtered, peptide levels. The significant differences in endogenous and exogenous peptide accumulation in the plasma and the kidney speak to the independent regulation of the systemic and intrarenal RAS and explain their decoupling in experimental models of hypertension.

In general, model simulations confirm that Losartan attenuates the rise in intrarenal Ang II levels during the development of hypertension via the “key point break-down effect” (blocking all positive feedback within the kidney), as was suggested by Xu et al (2009). However, we hypothesize that this is not the primary mechanism by which Losartan acts to normalize blood pressure. Instead, model simulations suggest that it is Losartan’s blockage of the main intracellular uptake path (AT1R binding and internalization) that contributes to the drug’s blood pressure regulating effects. Indeed, with this pathway still intact exogenous Ang II would continue to accumulate intrarenally, and more importantly, intracellularly as any infusion progressed, causing blood pressure to rise. This is not observed experimentally. Therefore, we hypothesize that it is actually the shift from intracellular to extracellular Ang II localization within the kidney that prevents blood pressure from rising under Losartan administration: Without being able to bind to or enter renal (or systemic) cells, Ang II cannot regulate sodium reabsorption, vessel tone, initiate secondary hormone secretion, or perform any signalling that would lead to increases in blood pressure. The same principals would likely hold for other ARBs.

### 4.1 Model limitations and future extensions

A key limitation of the present model was the necessary assumption that renal renin activity remains constant following Losartan administration. While previous modelling results were used to justify this choice (see section 2.2; renal tissue compartment), future work should focus on gathering experimental measurements of renal renin activity before and after Losartan treatment at different doses in both control and hypertensive rats. Additionally, the model does not account for the effects of Ang II, and therefore of Losartan and EXP3174, on renal hemodynamic function. Recall that intrarenal Ang II influences renal blood flow and glomerular filtration rate via various AT1R-dependent mechanisms. Indeed, afferent arteriole resistance, efferent arteriole resistance, the structure of the glomerular filtration barrier, and renal autoregulatory mechanisms such as tubuloglomerular feedback and the myogenic response are all affected by AT1R-bound Ang II, and therefore by Losartan administration (Xu et al, 2009; Ahmed and Layton, 2020; Denton et al, 2000; Toke and Meyer, 2001; Yang et al, 2011). Given that so many mechanisms are at play, the effect of Losartan on glomerular filtration rate is variable, often depending on whether blood pressure falls in or out of the renal autoregulatory range (Xu et al, 2009). As a result, including a direct link between Losartan and the current model’s renal hemodynamic parameters would be intractable. In future work, this could be overcome by incorporating the present model into Ahmed and Layton (2020)’s whole-body blood pressure regulation model, which already considers renal autoregulatory mechanisms and afferent/efferent arteriole resistance as variables. By simulating Losartan administration under different autoregulatory conditions, the mechanisms contributing to the drug’s variable effect on renal hemodynamics could be investigated.

The resulting Losartan-blood pressure regulation model could also be used to study how the shift from intracellular to extracellular intrarenal Ang II localization influences the stimulation of sodium reabsorption and consequently, mean arterial pressure (also variables in Ahmed and Layton (2020)’s model). To study Losartan’s effect on sodium reabsorption in more detail however, the present model could also be coupled to a kidney function model that stimulates epithelial transport of electrolytes and water (Layton and Layton, 2019; Layton et al, 2017; Sgouralis et al, 2017; Edwards and Layton, 2014; Chen et al, 2011). This coupling can be formulated based on the many known connections between Ang II, AT1Rs, and ion transport along the nephron (Burns and Li, 2003; Valles et al, 2005; Xu et al, 2009). Such a model could then be used to elucidate which transporters and nephron segments most greatly influence Losartan’s normalization of renal excretory function (Xu et al, 2009). The Losartan- and RAS-mediated electrolyte transport predicted by the kidney model could subsequently be used to modulate glomerular filtration rate via the tubuloglomerular feedback (Edwards et al, 2014; Layton, 2010), thereby influencing intrarenal RAS distribution.

It has been long established that males and females respond differently to anti-hypertensive therapies, including ARBs like Losartan (Sullivan, 2008; Hudson et al, 2007; Leete et al, 2018). The current model is limited in that it does not consider sex differences, and our results are therefore specific to a male rat’s response to Losartan treatment. This limitation could be overcome in future work by parameterizing sex-specific intrarenal and systemic model compartments using the procedure discussed in Smith and Layton (2023). The sex differences in the response of various RAS peptides to ARB treatment, which to our knowledge has not been well characterized in experiments, could then be simulated. These results could shed light on why male hypertensive rats tend to exhibit a greater reduction in blood pressure following Losartan administration than females (Yanes et al, 2006). The sex-specific PK models could also be coupled to a sex-specific blood pressure regulation model (Ahmed and Layton, 2020), a sex-specific epithelial solute transport model (Li et al, 2018; Hu et al, 2019, 2020, 2021), or a sex-specific renal blood flow model (Chen et al, 2017) to study the sex-specific effects of Losartan on blood pressure, sodium reabsorption, and renal hemodynamics in more detail, respectively.

Another natural extension of this work would be the creation of a PK model of a common ACE inhibitor (ACEi). In this way, the effects of the two classes of anti-hypertensive therapies on the intrarenal RAS could be compared. Coupling can be achieved through systemic and renal ACEi–based inhibition of ACE activity 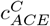 in all model compartments *C*. We hypothesize that ACEis, like ARBs, will prevent Ang II–induced hypertension, at least in part, by blocking the AT1R–mediated accumulation of Ang II in tubular epithelial cells. However, this class of drug will achieve this via a different mechanism: ACEi will reduce the endogenous production of Ang II in the kidney and thus, the pool of Ang II that can be internalized. We expect ARBs to be more effective than ACEis at preventing hypertension induced by Ang II infusion because ACEis would not prevent the accumulation of exogenous Ang II in tubular epithelial cells following Ang II infusion, but ARBs do (as demonstrated above).

Finally, our model does not represent the circadian rhythms known to be present in the systemic RAS (Hilfenhaus, 1976), and likely to be present in the intrarenal RAS. To explore whether the timing of Losartan administration influences its effectiveness as an anti-hypertensive therapy, the model could be extended to include circadian variations in appropriate model parameters. The resulting model could also be coupled to a kidney function model that represents circadian rhythms (Layton and Gumz, 2022; Wei et al, 2018) to study whether the daily variations in RAS peptide levels impact volume homeostasis.

## 5 Conclusion

In this work, we discussed the varying effects of ARBs on the intrarenal and systemic RASs in hypertension induced by Ang II infusion. We found that ARBs more effectively inhibit the activity of the intrarenal RAS due to the nature of it’s independent regulatory mechanisms, particularly *UPTK*. Since this mechanism also facilitates the decoupling of the two systems, a signature that is common to many experimental models of hypertension that are more representative of clinical hypertension, the conclusions drawn from this work are likely to have broader implications to cases of clinical hypertension that are associated with an over-active RAS. Our hypothesis, that *the intrarenal RAS is the primary pharmacological target of ARB treatment*, can be further explored in future work by extending the present model to study these other experimental models of hypertension that do not involve the constant infusion of unphysiological doses of Ang II, such as: two-kidney, one-clip Goldblatt hypertension Cervenka et al (1999), salt-sensitive rats Wu et al (2014), and spontaneously hypertensive rats Takenaka et al (1990).

## Statements and Declarations

Funding: This work was supported by the Canada 150 Research Chair program and by the Natural Sciences and Engineering Research Council of Canada, via a Discovery Award (to Anita Layton) and a Canadian Graduate Scholarship (to Delaney Smith).

- Competing interests: None.
- Ethics approval: Not applicable.
- Consent to participate: Not applicable.
- Consent for publication: Not applicable.
- Availability of data and materials: Not applicable.
- Code availability: https://github.com/Layton-Lab/intrarenalRAS.git
- Authors’ contributions: Conceptualization: Delaney Smith, Anita Layton; Methodology: Delaney Smith; Formal Analysis and Investigation: Delaney Smith; Writing (original draft preparation): Delaney Smith; Writing (review and editing): Delaney Smith, Anita Layton; Supervision: Anita Layton.

Editorial Policies for:

Springer journals and proceedings:

https://www.springer.com/gp/editorial-policies

Nature Portfolio journals:

https://www.nature.com/nature-research/editorial-policies

*Scientific Reports*:

https://www.nature.com/srep/journal-policies/editorial-policies

BMC journals:

https://www.biomedcentral.com/getpublished/editorial-policies

## Appendix A Intrarenal Losartan-EXP3174 model equations

Below, the model equations describing the dynamics of metabolite *X* (*X* = *Los* or *EXP* 3174) in each extracellular intrarenal sub-compartment *C* (*C* = *Gl, Tb, Pt*, or *Pv*) are summarized.

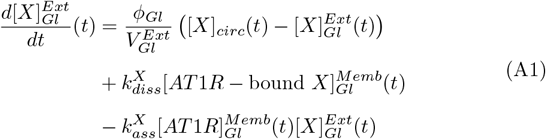

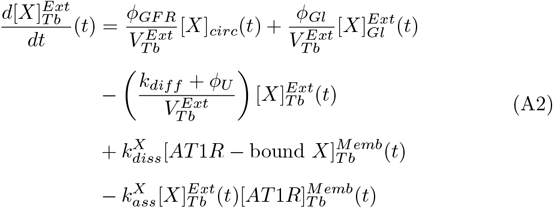

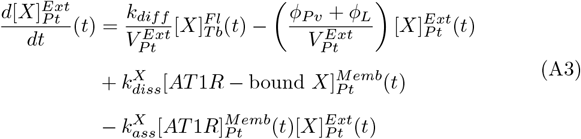

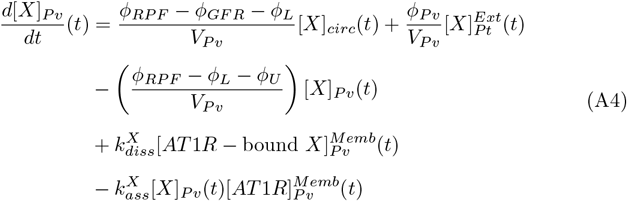

